## Supplementary figures and images for "Multi-epitope immunocapture of huntingtin reveals striatum-selective molecular signatures"

### Figure S1

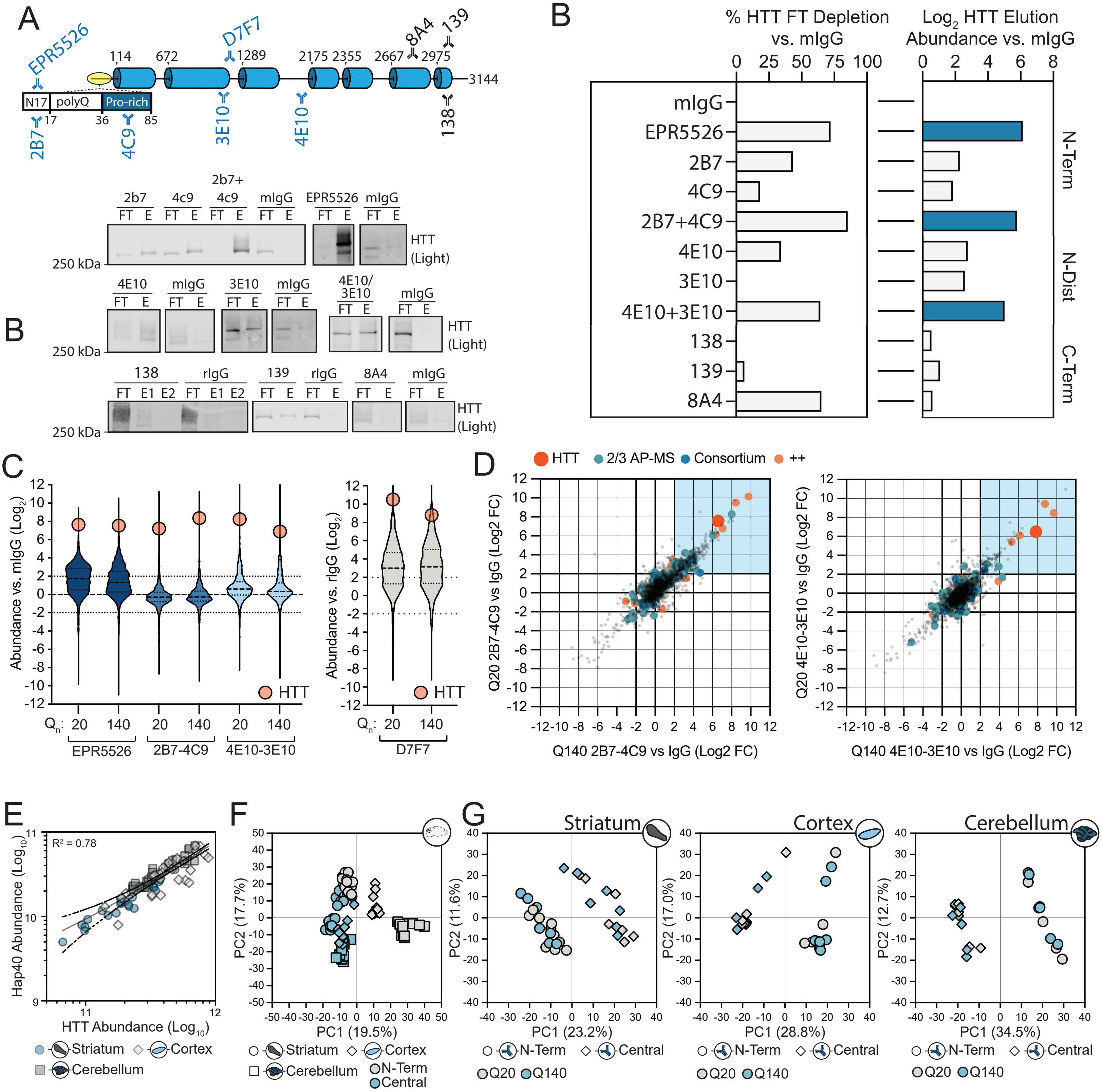

### Figure S2

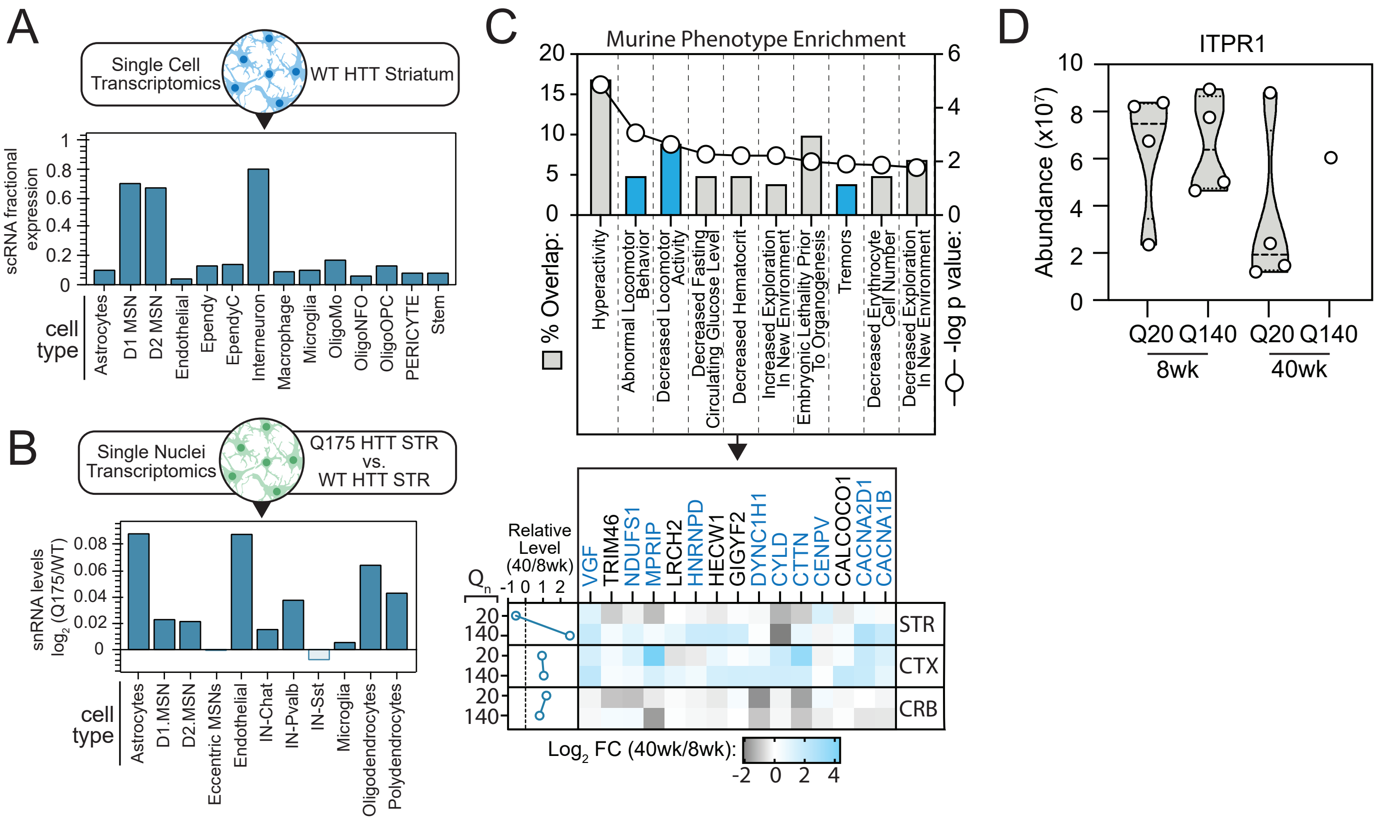
